# Targeting ALCAT1 Ameliorates Cardiomyopathy in Barth Syndrome

**DOI:** 10.64898/2026.09.24.754267

**Authors:** Jun Zhang, Qianqian Ye, Xi Fang, Wendong Huang, Yuguang Shi

## Abstract

**Background:** Barth syndrome (BTHS) is an X-linked mitochondrial disorder caused by loss-of-function mutations in *TAFAZZIN* (*TAZ*), resulting in defective cardiolipin (CL) remodeling, cardiomyopathy, and premature death. The mechanisms linking *TAZ* deficiency to impaired mitochondrial quality control remain incompletely understood, and there are no disease-modifying therapies available for BTHS. Here, we investigated the role of Acyl-CoA Lysocardiolipin Acyltransferase-1 (ALCAT1), a stress-inducible phospholipid-remodeling enzyme, in BTHS cardiomyopathy.

**Methods:** Inducible *TAZ* knockdown (*TAZ_KD_*) and cardiomyocyte-specific *TAZ* knockout (*TAZ^cKO^*) mice were used to determine the effects of genetic *ALCAT1* deletion or pharmacological inhibition with Juvenatin, a highly selective small molecule ALCAT1 inhibitor. Cardiac function, exercise capacity, mitochondrial function, lipid remodeling, lysosomal function, and mitophagic flux were assessed.

**Results:** ALCAT1 was markedly upregulated in *TAZ*-deficient hearts. Genetic *ALCAT1* deletion attenuated cardiac dysfunction in *TAZ_KD_* mice. Juvenatin similarly improved cardiac function in *TAZ_KD_* mice and reversed established cardiomyopathy in *TAZ^cKO^* mice. Genetic or pharmacological ALCAT1 inhibition improved mitochondrial ultrastructure and respiration and reduced mitochondrial oxidative stress. These benefits occurred without correcting the characteristic CL abnormalities caused by *TAZ* deficiency. Instead, ALCAT1 deletion or inhibition corrected pathological phosphatidylinositol remodeling, normalized phosphoinositide signaling, and restored lysosomal function and mitophagic flux, leading to improved mitochondrial quality control in *TAZ*-deficient cells.

**Conclusions:** ALCAT1 contributes to BTHS cardiomyopathy through a CL-independent pathway involving dysregulated phosphatidylinositol/phosphoinositide signaling, lysosomal dysfunction, and impaired mitophagy. Genetic and pharmacological targeting of ALCAT1 restores mitochondrial and cardiac function despite persistent CL abnormalities, establishing ALCAT1 as a promising disease-modifying therapeutic target for BTHS.

## Introduction

Barth syndrome (BTHS) is a lethal, X-linked genetic disorder characterized by dilated cardiomyopathy, skeletal myopathy, neutropenia, and growth retardation, predominantly found in males^1,2^. BTHS is caused by mutations in *TAFAZZIN* (*TAZ*) gene, which encodes a mitochondrial phospholipid transacylase localized to membranes facing the intermembrane space^3,4^. TAZ is required for the remodeling of cardiolipin (CL), a signature mitochondrial phospholipid that plays critical roles in maintaining mitochondrial membrane architecture, cristae organization, respiratory chain assembly, and bioenergetic function^5,6^. Loss-of-function mutations in the *TAZ* gene result in depletion of tetralinoleoyl CL (TLCL), the predominant and functionally optimal CL specie in the heart, liver, and skeletal muscle^4,7^. Non-functional TAZ results in TLCL deficiency and accumulation of monolysocardiolipin (MLCL), a central driver of mitochondrial dysfunction, cardiomyopathy, neutropenia, and growth defects in BTHS^1,8^. However, the downstream molecular mechanisms linking CL defects to cardiac dysfunction remain incompletely understood. Although supportive care has improved survival, there are no disease-effective therapies for BTHS, highlighting the need to identify new therapeutic targets and mechanisms underlying disease progression.

Despite the central role of CL in mitochondrial biology, increasing evidence suggests that other mechanisms contribute to disease progression in BTHS. TAZ deficiency was linked to abnormalities in mitochondrial dynamics, oxidative phosphorylation, reactive oxygen species (ROS) production, and mitochondrial quality control pathways^9–13^. In particular, defective mitophagy, a selective autophagy of mitochondria that plays a critical role in mitochondrial quality control and cell survival by clearing damaged mitochondria, participates in BTHS cardiomyopathy^10,11,13–16^. Restoration of mitophagy mitigated mitochondrial dysfunction and cardiomyopathy in a mouse model of BTHS^14^. However, the molecular mechanisms linking TAZ deficiency to impaired mitophagy in BTHS remain poorly defined.

Acyl-CoA:lysocardiolipin acyltransferase-1 (ALCAT1) is a stress-inducible acyltransferase that catalyzes CL remodeling by incorporating polyunsaturated fatty acids into CL, generating species that are highly susceptible to peroxidation^17,18^. Upregulated ALCAT1 protein expression by oxidative stress induces mitochondrial ROS production, mtDNA instability and mitochondrial dysfunction, and is implicated in the mitochondrial etiology of various metabolic disorders^19–21^. Consequently, genetic depletion or pharmacological inhibition of the ALCAT1 in mice restored mitochondrial function and mitigated age-related metabolic diseases^18–20,22–27^. In addition to CL remodeling, ALCAT1 acts as a lysophosphatidylinositol acyltransferase to catalyze the remodeling of phosphatidylinositol (PI)^28^, the precursor for phosphoinositides (PIPs), a family of signaling lipids that regulate membrane trafficking, autophagy, lysosomal function, and mitochondrial quality control^29^. In this study, we tested the hypothesis that ALCAT1 amplifies mitochondrial stress and disrupts mitochondrial quality control to promote BTHS. Using genetic and pharmacological approaches in inducible TAZ knockdown and cardiac-specific TAZ knockout mouse models, we demonstrate that ALCAT1 is strongly upregulated in TAZ-deficient hearts. Moreover, genetic ablation or pharmacological inhibition with the potent selective small-molecule inhibitor Juvenatin (Juve), attenuate dilated cardiomyopathy and left ventricular (LV) dysfunction in mouse models of BTHS. Mechanistically, ALCAT1 inhibition did not correct CL abnormalities but instead normalized maladaptive PI remodeling and PIPs accumulation, leading to restoration of lysosomal function and re-established mitophagic flux in TAZ-deficient cells. These findings uncover a previously unrecognized ALCAT1-PI-PIP signaling axis that links TAZ deficiency to impaired mitochondrial quality control. The results also implicate Juve, a first-in-class ALCAT1 inhibitor, as a promising therapeutic compound for BTHS.

## Results

### ALCAT1 deficiency attenuates cardiomyopathy and LV dysfunction in TAZ knockdown mice

ALCAT1 and TAZ are two key enzymes in CL remodeling. We found that ablation of ALCAT1 significantly increased cardiac TAZ expression (Figure 1A and 1B), while TAZ deficiency markedly increased cardiac ALCAT1 protein levels (Figure 1C and 1D), indicating that these two enzymes appear to be intricately interconnected and a potential pathogenic role of ALCAT1 in BTHS. To further interrogate the role of ALCAT1 in BTHS, whole-body ALCAT1 knockout mice^18^ were mated with the inducible *TAZ* knockdown (TAZ_KD_) transgenic mice^30^ to generate ALCAT1 knockout mice with deficient TAZ expression (KO/KD). Wild-type (WT), TAZ_KD_, and KO/KD mice were fed a doxycycline-containing diet (625 mg/kg) beginning at one month of age for seven months (Figure 1E). TAZ was efficiently decreased in the hearts of both TAZ_KD_ and KO/KD mice (Figure 1C and 1D). Consistent with the precious report^30^, TAZ_KD_ mice exhibited impaired growth (Figure 1F), and severe dilated cardiomyopathy and LV dysfunction. Echocardiography showed enlarged LV internal dimensions in systole (LVIDs) and reduction in LV ejection fraction (EF) and fractional shortening (FS) compared with control mice (Figure 1G to 1J). ALCAT1 deficiency rescued the growth delay (Figure 1F) and attenuated cardiac abnormalities, with normalization of systolic LVID and increased LV EF and FS relative to TAZ_KD_ mice (Figure 1G to 1J). TAZ deficiency also caused myocardial hypertrophy, as demonstrated by enlarged cardiomyocytes in hematoxylin and eosin (H&E) staining (Figure 1K and 1L). In contrast, ablation of ALCAT1 effectively prevented myocardial hypertrophy, as shown by decreased cardiomyocyte size (Figure 1K and 1L). Consistent with these findings, ALCAT1 deficiency also significantly decreased mRNA expression levels of genes associated with cardiac hypertrophy, including atrial natriuretic factor (*Anf*), β-cardiac myosin heavy chain (*β-Mhc*), brain natriuretic peptide (*Bnp*), and alpha-skeletal actin (*Acta1*), in the heart of TAZ_KD_ mice (Figure 1M). Ablation of ALCAT1 significantly retarded the accumulation of cardiac collagen, as shown by Masson’s trichrome staining of LV samples (Figure 1N and 1O; arrows highlight collagen fibers). Furthermore, ablation of ALCAT1 also decreased the mRNA expression levels of pro-fibrotic genes, including *Collagen 1* and *Collagen 3*, in the heart of TAZ_KD_ mice (Figure 1M).

**Figure 1.**
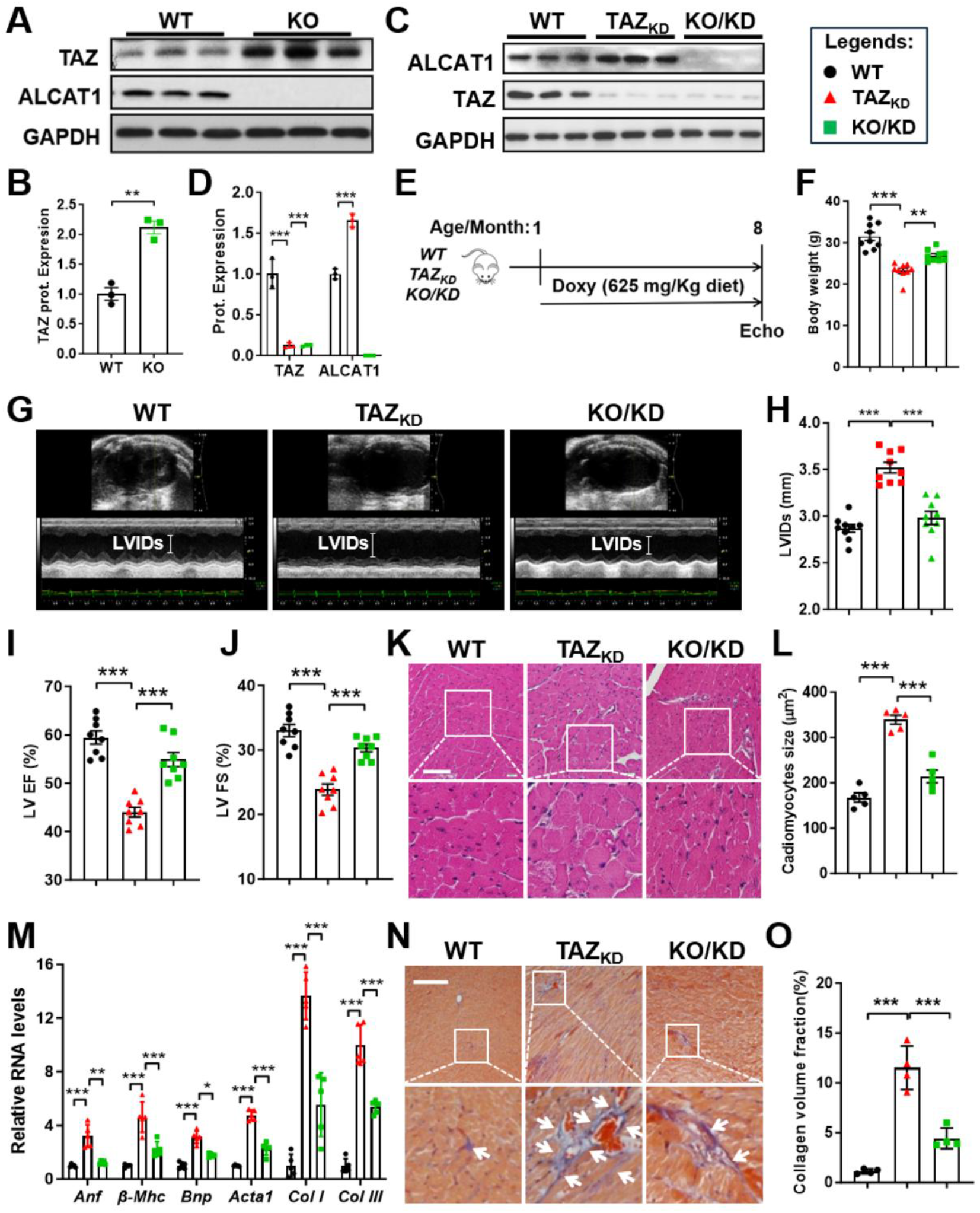
ALCAT1 deficiency attenuates cardiomyopathy and LV dysfunction in TAZ_KD_ mice. **A-B**, Western blot (**A**) and quantitative analysis (**B**) of ALCAT1 and TAZ in WT and ALCAT1 KO murine hearts. n = 3. **C**-**D**, Western blot (**C**) and quantitative analysis (**D**) of ALCAT1 and TAZ in WT, TAZ_KD_, and KO/KD murine hearts. n = 3. **E**, Experimental outline. WT, TAZ_KD_, and KO/KD mice were fed a diet containing doxycycline (625 mg/Kg) starting at 1-month of age for 7 consecutive months. **F**, Body weight WT, TAZ_KD_, and KO/KD mice at 8-months of age. n = 9. **G**-**J**, Representative echocardiographic images (**G**), LV internal dimensions at systole (LVIDs, **H**), LV ejection fraction (**I**), and fractional shortening (**J**) of WT, TAZ_KD_, and KO/KD mice at 8-months of age. n=8. **K**-**L**, H&E staining of LV sections (**K**) and quantification analysis of cardiomyocytes size (**L**) in WT, TAZ_KD_, and KO/KD mice. n = 5. Scale bar, 50 μm. **M**, qPCR analysis of mRNA expression levels of genes associated with cardiac hypertrophy and fibrosis in WT, TAZ_KD_, and KO/KD mice hearts. n=5. **N**-**O**, Masson’s trichrome staining of collagen fibers (**N**) and quantitative analysis of collagen volume fraction (**O**) in WT, TAZ_KD_, and KO/KD murine heart tissue sections. Arrows highlight collagen fibers. n=5. Scale bar, 50 μm. Data are presented as mean ± SEM. Statistical analysis was performed using one-way ANOVA with Tukey’s post hoc test or unpaired Student t-test. **p<0.01, ***p<0.001.

### Pharmacological inhibition of ALCAT1 ameliorates cardiomyopathy and functional impairment in mouse models of BTHS

To evaluate the therapeutic potential of targeting ALCAT1 in BTHS, we treated the TAZ_KD_ mice with Juve (previously named Dafaglitapin), a highly potent and selective small-molecule inhibitor of ALCAT1^24^. Mice were given doxycycline-containing water (2 mg/mL) starting at 1 month of age to induce TAZ knockdown, followed by oral gavage of vehicle (5% Carboxymethylcellulose) or Juve (10 mg/kg body weight) daily from 2 to 8 months of age (Figure 2A). Compared with the vehicle-treated mice, inhibition of ALCAT1 by Juve significantly mitigated body weight loss associated with TAZ knockdown (Figure 2B). Quantitative magnetic resonance imaging analysis demonstrated that Juve treatment increased lean body mass, with no meaningful effect on fat mass in TAZ_KD_ mice (Figure 2C). Consistently, Juve treated animals showed improved grip strength, which was reduced in TAZ_KD_ mice (Figure 2D). Moreover, TAZ deficiency resulted in reduced treadmill running distance and work output, both of which were substantially restored after treatment with Juve (Figure 2E). Inhibition of ALCAT1 with Juve also prevented dilated cardiomyopathy and restored LV function (Figure 2F), as shown by reduced LVID at both systole and diastole (Figure 2G and 2H), and increased interventricular septal wall thickness and LV posterior wall thickness at systole (Figure 2I and 2J), resulting in increased LV EF and FS (Figure 2K and 2L). This was associated with less myocardial fibrosis in TAZ_KD_ mice, as demonstrated by results from Masson’s trichrome staining of LV samples (Figure 2M and 2N) and downregulated mRNA expression levels of *Collagen 1* and *Collagen 3* in the heart of TAZ_KD_ mice (Figure 2O).

**Figure 2.**
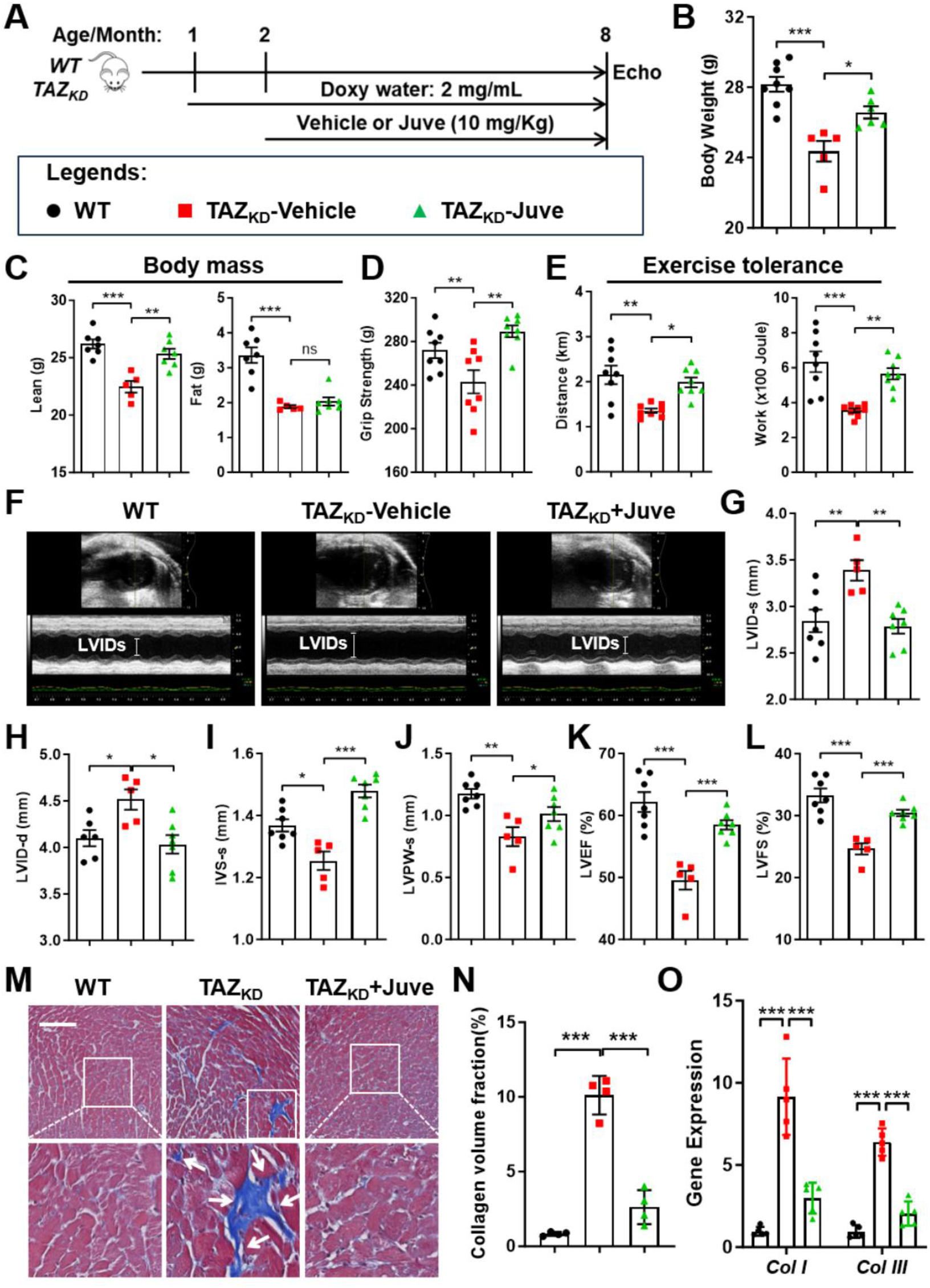
Pharmacological inhibition of ALCAT1 ameliorates cardiomyopathy and functional impairment in TAZ_KD_ mice. **A**, Experimental outline. WT and TAZ_KD_ mice had unrestricted access to water containing doxycycline (2 mg/mL) starting at 1-month of age to knockdown TAZ in the transgenic mice. Mice were then oral gavaged daily with vehicle (5% Carboxymethylcellulose) or Juve (10 mg/Kg BW) starting at 2-months of age for 6 months. **B**-**C**, Body weight (**B**) and qMRI analysis of body mass (**C**) of WT and TAZ_KD_ mice treated with vehicle or Juve. n = 5-8. **D**-**E**, Grip strength test (**D**) and treadmill exercise tolerance test (**E**) in WT and TAZ_KD_ mice treated with vehicle or Juve. n = 8. **F**-**L**, Representative echocardiographic images (**F**), LVIDs (**G**), LVIDd (**H**), IVSs (**I**), LVPWs (**J**), LV EF (**K**), and FS (**L**) of WT and TAZ_KD_ mice treated with vehicle or Juve. n=5-7. **M**-**N**, Masson’s trichrome staining of collagen fibers (**M**) and quantitative analysis of collagen volume fraction (**N**) in heart tissue sections of WT and TAZ_KD_ mice treated with vehicle or Juve. Arrows highlight collagen fibers. n=4. Scale bar, 50 μm. **O**, qPCR analysis of mRNA expression levels of *Collagen I* and *Collagen III* in the hearts of WT and TAZ_KD_ mice treated with vehicle or Juve. n=5. Data are presented as mean ± SEM. Statistical analysis was performed using one-way ANOVA with Tukey’s post hoc test. *p<0.01, **p<0.01, ***p<0.001.

The TAZ_KD_ mice developed dilated cardiomyopathy at much older age due to incomplete inactivation of TAZ^31^. Closer to clinical BTHS, we treated cardiac-specific TAZ knockout (TAZ^cKO^) mice with Juve. The TAZ^cKO^ mice, which were generated by crossing the *TAZ* floxed mice^12^ with the *Myh6-Cre* mice, developed cardiomyopathy at ∼2 months of age, as others reported^12,32^. Beginning at 3-months of age, the TAZ^cKO^ mice were fed a control diet or diet supplemented with Juve (150 mg/Kg) for 3 consecutive months (Figure 3A). Juve treatment significantly attenuated dilated cardiomyopathy and restored LV function in TAZ^cKO^ mice (Figure 3B), and reduced LVIDs and increased IVSs, LVPWs and LVPWd (Figure 3C to 3F), resulting in recovery of LV EF and FS (Figure 3G and 3H). Juve treatment also improved exercise capacity of TAZ^cKO^ mice, as evidenced by the increased running distance and work output in treadmill test (Figure 3I). In addition, TAZ^cKO^ mice treated with Juve had decreased cardiomyocyte size (Figure 3J and 3K) and downregulated the levels of RNAs associated with cardiac hypertrophy (Figure 3L). Moreover, Juve treatment also reduced collagen deposition in cardiac muscle (Figure 3M and 3N).

**Figure 3.**
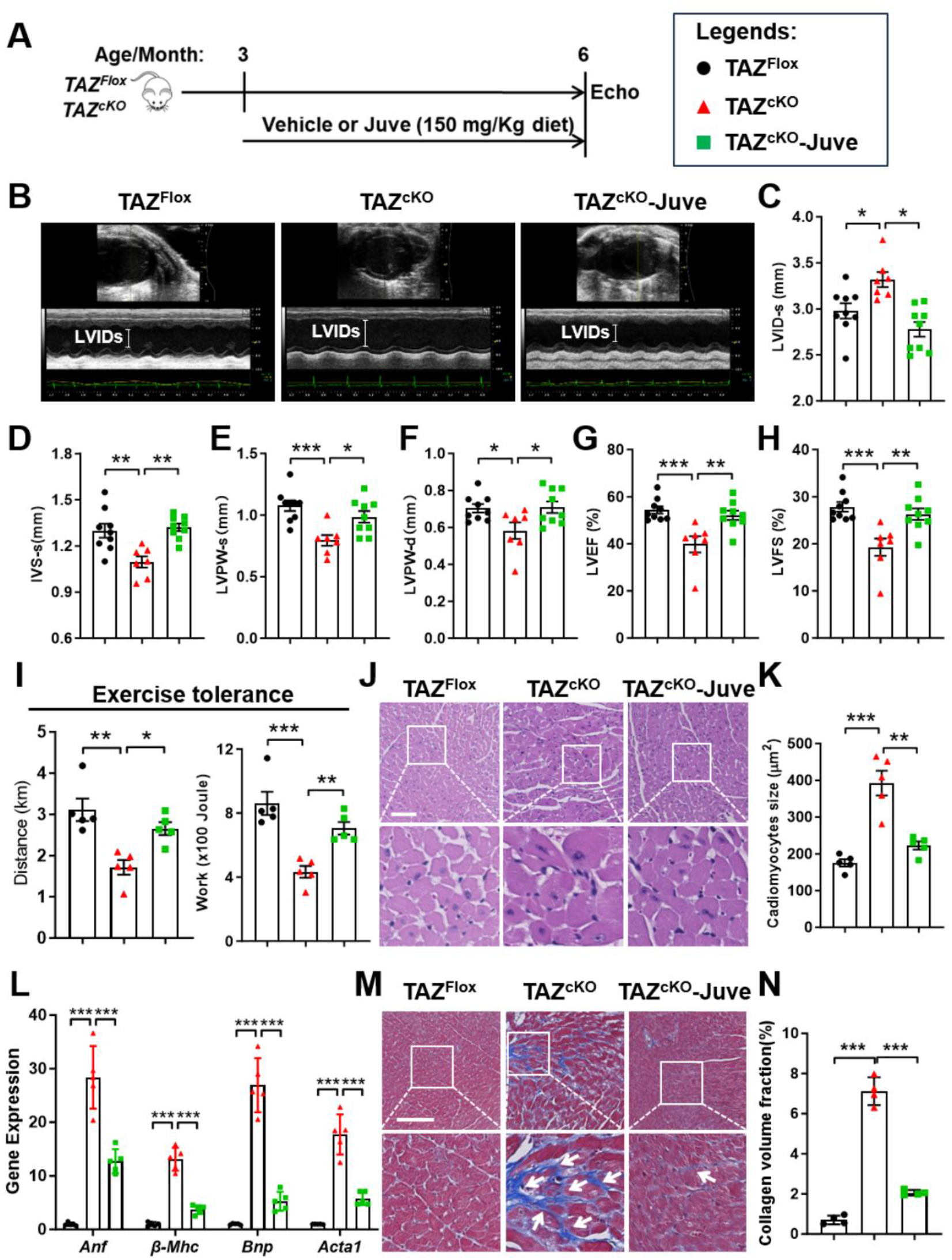
Pharmacological inhibition of ALCAT1 mitigates established cardiomyopathy in TAZ^cKO^ mice. **A**, Experimental outline. TAZ floxed mice were bred with *Myh6-Cre* mice to generate TAZ cardiomyocyte-specific KO (TAZ^cKO^) mice. TAZ floxed and TAZ^cKO^ mice were fed a control diet or a diet containing Juve (150 mg/Kg diet) starting at 3-months of age for 3 consecutive months. **B**-**H**, Representative echocardiographic images (**B**), LVIDs (**C**), IVSs (**D**), LVPWs (**E**), LVPWd (**F**), LV EF (**G**, and FS (**H**) of TAZ floxed and TAZ^cKO^ mice treated with vehicle or Juve. n=7-9. **I**, Treadmill exercise tolerance test results of TAZ floxed and TAZ^cKO^ mice treated with vehicle or Juve. n = 5. **J**-**K**, H&E staining of LV sections (**J**) and quantification analysis of cardiomyocytes size (**K**) in TAZ floxed and TAZ^cKO^ mice treated with vehicle or Juve. n = 5. Scale bar, 50 μm. **L**, qPCR analysis of mRNA expression levels of genes associated with cardiac hypertrophy in the hearts of TAZ floxed and TAZ^cKO^ mice treated with vehicle or Juve. n=5. **M**-**N**, Masson’s trichrome staining of collagen fibers (**M**) and quantitative analysis of collagen volume fraction (**N**) in the heart tissue section from TAZ floxed and TAZ^cKO^ mice treated with vehicle or Juve. Arrows highlight collagen fibers. n=4. Scale bar, 50 μm. Data are presented as mean ± SEM. Statistical analysis was performed using one-way ANOVA with Tukey’s post hoc test. *p<0.05, **p<0.01, ***p<0.001.

### ALCAT1 inhibition restores mitochondrial integrity and function in TAZ-deficient cardiomyocytes

One of the major cellular defects in cells from individuals with BTHS is the accumulation of structurally abnormal mitochondria due to defective mitochondrial quality control. Transmission electron microscopy (TEM) of LV sections from TAZ_KD_ mice revealed abnormalities in mitochondrial morphology, including fragmentation, irregular shape and distribution, and disrupted cristae architecture (Figure 4A and 4B). Notably, TAZ deficiency also led to accumulation of autophagic vacuoles wrapped by mitochondria (Figure 4A and 4C), indicating defects in mitophagy. Ablation of ALCAT1 improved mitochondrial morphology, restored cristae organization, and significantly reduced the accumulation of autophagic vacuoles in TAZ-deficient cardiomyocytes (Figure 4A to 4C), suggesting normalization of mitochondrial turnover. Consistent with these structural improvements, ALCAT1 deficiency normalized mitochondrial DNA (mtDNA) copy number which was increased in TAZ-deficient cardiomyocytes (Figure 4D), and attenuated cardiac mitochondrial ROS in the hearts of TAZ_KD_ mice (Figure 4E). To determine if the morphologic improvements had functional implications, we analyzed mitochondrial respiration using Seahorse extracellular flux analysis. Cardiomyocytes isolated from TAZ_KD_ mice had reductions in basal respiration, ATP-linked respiration, maximal respiratory capacity, and spare respiratory capacity (Figure 4F and 4G). Deletion of ALCAT1 significantly restored the respiratory parameters in TAZ_KD_ cardiomyocytes (Figure 4F and 4G), indicating a recovery of mitochondrial function.

**Figure 4.**
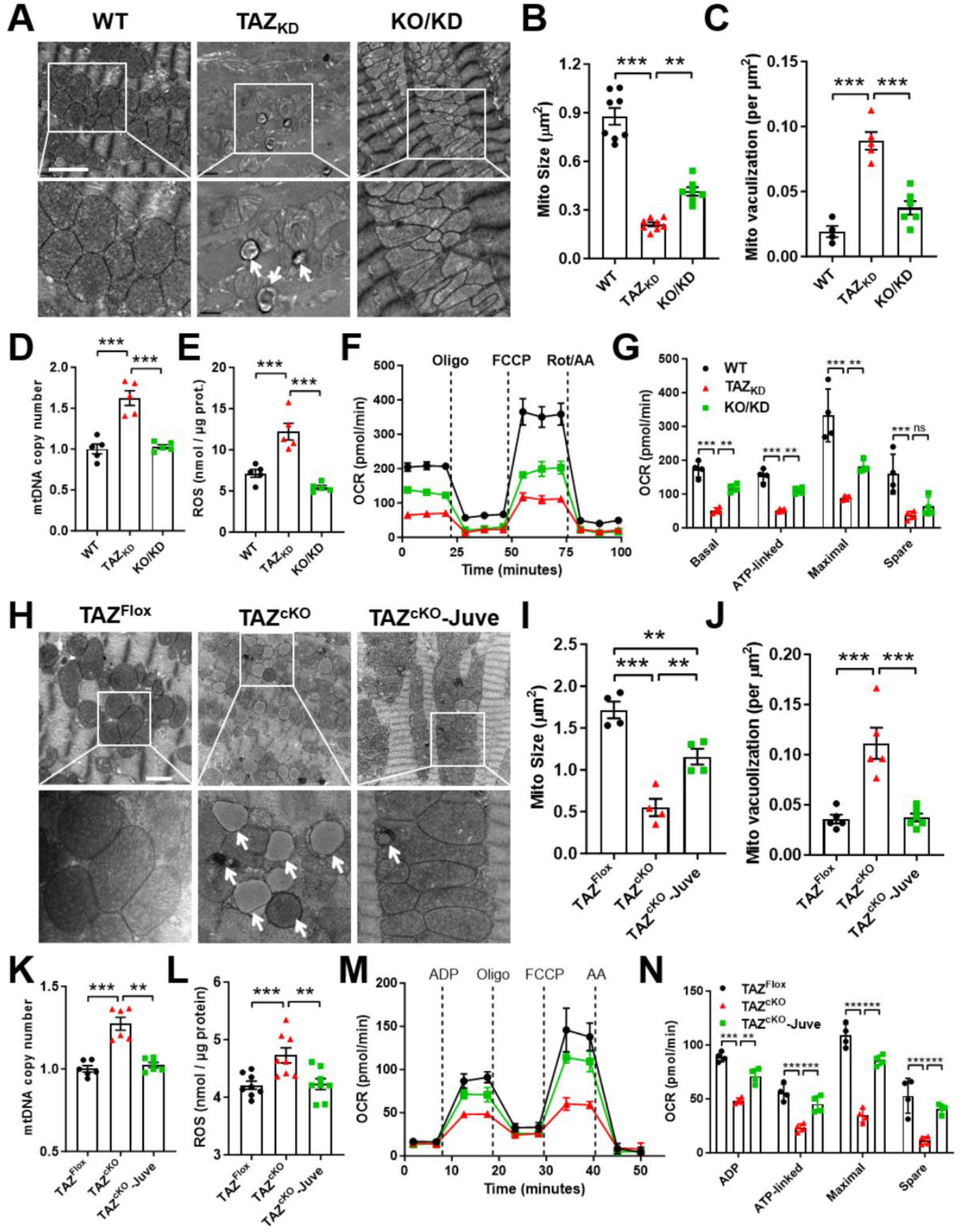
ALCAT1 deficiency or inhibition restores mitochondrial integrity and function in TAZ-deficient cardiomyocytes. **A-C**, TEM analysis of mitochondria morphology (**A**) and quantitative analysis of mitochondrial size (**B**) and mitochondrial vacuolization rate (**C**) in the hearts of WT, TAZ_KD_, and KO/KD mice. n = 6-8. Scale bar, 2 μm. **D**, qPCR analysis of mtDNA copy number in the hearts of WT, TAZ_KD_, and KO/KD mice. n = 5. **E**, ROS production measurement in isolated mitochondria from WT, TAZ_KD_, and KO/KD murine hearts. n = 5. **F**-**G**, Seahorse analysis (**F**) and quantitative analysis (**G**) of mitochondrial oxygen consumption rate (OCR) in cardiomyocytes isolated from WT, TAZ_KD_, and KO/KD murine hearts. n = 5. **H**-**J**, TEM analysis of mitochondria morphology (**H**) and quantitative analysis of mitochondrial size (**I**) and vacuolization rate (**J**) in the hearts of TAZ floxed and TAZ^cKO^ mice treated with vehicle or Juve. n = 4. Scale bar, 2 μm. **K**, qPCR analysis of mtDNA copy number in the hearts of TAZ floxed and TAZ^cKO^ mice treated with vehicle or Juve. n = 5. **L**, ROS production in isolated mitochondria from the hearts of TAZ floxed and TAZ^cKO^ mice treated with vehicle or Juve. n = 8. **M**-**N**, Seahorse analysis (**M**) and quantitative analysis (**N**) of mitochondrial OCR in isolated mitochondria from the hearts of TAZ floxed and TAZ^cKO^ mice treated with vehicle or Juve. n = 4. Data are presented as mean ± SEM. Statistical analysis was performed using one-way ANOVA with Tukey’s post hoc test. **p<0.01, ***p<0.001.

To determine whether pharmacological inhibition of ALCAT1 produced similar mitochondrial benefits, we analyzed mitochondrial morphology and function in the heart of Juve-treated TAZ_KD_ and TAZ^cKO^ mice. Juve treatment restored mitochondrial ultrastructure and reduced the number of autophagic vacuoles in both TAZ^cKO^ and TAZ_KD_ mice, as determined by TEM analysis of LV sections (Figure 4H to 4J, and Figure S1A to S1C). Juve treatment also normalized mtDNA copy number (Figure 4K) and reduced mitochondrial ROS production (Figure 4L, and Figure S1D) in TAZ-deficient cardiomyocytes. Moreover, Juve restored mitochondrial respiration, as evidenced by Seahorse analysis in mitochondria isolated from the hearts of TAZ^cKO^ (Figure 4M and 4N) and TAZ_KD_ mice (Figure S1E and S1F).

### ALCAT1 deficiency or inhibition mitigates defective PI remodeling and PIPs accumulation in TAZ-deficient hearts

To elucidate the molecular mechanisms that might account for how ALCAT1 regulates mitochondrial function in BTHS, we performed comprehensive lipidomic profiling in the hearts from both TAZ-deficient mouse models. Since ALCAT1 catalyzes the pathological remodeling of CL^17,18,20^, we examined the changes of CL profile by ALCAT1 deficiency or inhibition in both TAZ_KD_ and TAZ^cKO^ mice hearts. As expected, TAZ deficiency significantly decreased total CL and TLCL with increased MLCL, a hallmark of CL remodeling defects in BTHS, in both TAZ_KD_ and TAZ^cKO^ hearts (Figure S2A to S2F). Neither ALCAT1 genetic deletion nor pharmacological inhibition by Juve restored the CL abnormalities (Figure S2A to S2F). These results indicate that the cardioprotective effects of ALCAT1 inhibition are not mediated through restoration of the primary CL defect, pointing toward a potential alternative lipid pathway.

Beyond its role in CL remodeling, ALCAT1 also catalyzes the remodeling of PI^28^, a precursor for PIPs which are essential regulators of autophagy, mitophagy, membrane trafficking, and lysosomal signaling^33–35^. We examined whether ALCAT1 modulates the PI-PIP axis in TAZ-deficient hearts. Lipidomic analysis revealed that TAZ deficiency significantly elevated total PI levels and the abundance of the predominant PI species, PI-38:4, in the heart of the mutant mice (Figure 5A to 5D). ALCAT1 deficiency or inhibition with Juve significantly normalized both total PI and PI-38:4 in the TAZ-deficient hearts versus WT controls (Figure 5A to 5D). TAZ deficiency also significantly increased the levels of downstream PIP species, including phosphatidylinositol phosphate (PIP1), phosphatidylinositol bisphosphate (PIP2), and phosphatidylinositol trisphosphate (PIP3), in the hearts of TAZ^cKO^ mice (Figure 5E to 5J). Again, Juve treatment reduced the abundance of all three PIP species to the levels of the WT mice (Figure 5E to 5J). These results demonstrate that upregulated ALCAT1 expression contributes to aberrant PI remodeling and PIP accumulation in TAZ-deficient cardiomyocytes, suggesting that correction of PI remodeling, rather than restoration of CL, by ALCAT1 inhibition leads to restoration of mitochondrial quality control and cardiac function in BTHS.

**Figure 5.**
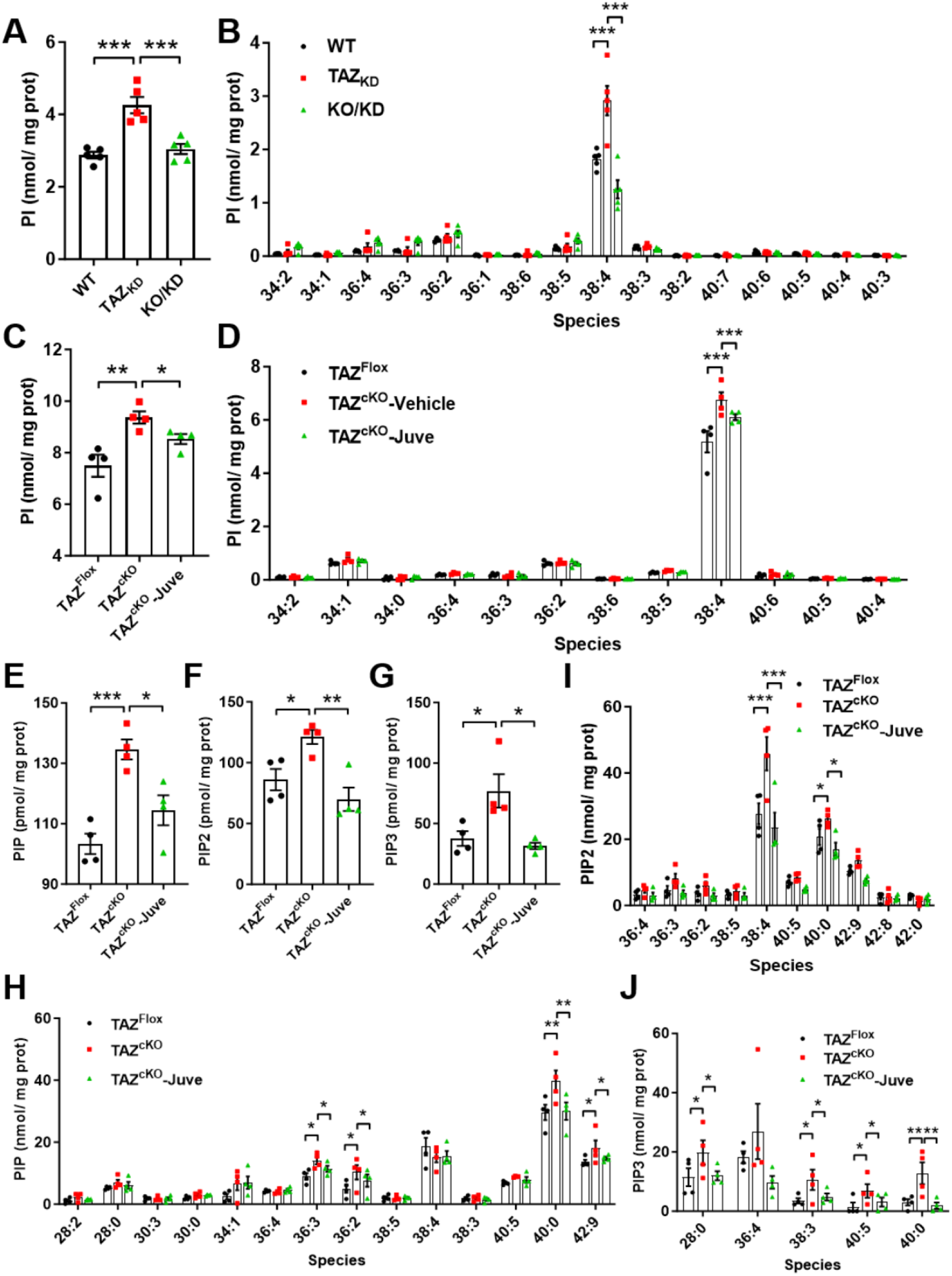
ALCAT1 deficiency or inhibition mitigates defective PI remodeling and PIPs accumulation in TAZ-deficient hearts. **A-B**, Lipidomic analysis of PI (**A**) and PI species (**B**) in the hearts of WT, TAZ_KD_, and KO/KD mice. n = 5. **C**-**D**, Lipidomic analysis of PI (**C**) and PI species (**D**) in the hearts of TAZ floxed and TAZ^cKO^ mice treated with vehicle or Juve. n = 4. **E**-**J**, Lipidomic analysis of PIP (**E**), PIP2 (**F**), PIP3 (**G**), PIP species (**H**), PIP2 species (**I**), and PIP3 species (**J**) in the hearts of TAZ floxed and TAZ^cKO^ mice treated with vehicle or Juve. n = 4. Data are presented as mean ± SEM. Statistical analysis was performed using one-way ANOVA with Tukey’s post hoc test. *p<0.05, **p<0.01, ***p<0.001.

### ALCAT1 deficiency or inhibition restores lysosome function and mitophagic flux in TAZ-deficient cells

The accumulation of mitochondria-containing autophagic vacuoles, together with altered levels of PIPs, suggested defective mitophagic flux in TAZ-deficient cardiomyocytes. Pursuing this we found that TAZ deficiency upregulated autophagy- and mitophagy-related proteins, including the LC3-I/LC3-II ratio, autophagy receptor protein p62, PARKIN and PINK1, in the hearts of TAZ_KD_ (Figure 6A, Figure S1G and S1H, and Figure S3A) and TAZ^cKO^ mice (Figure 6B and Figure S3B). Consistent with the reduced number of autophagic vacuoles, ablation of ALCAT1 or inhibition by Juve downregulated the expression of these biomarkers (Figure 6A and 6B, Figure S1G and S1H, and Figure S3), suggesting a restoration of autophagic consumption in TAZ-deficient cells. TAZ deficiency also induced the expression levels of class III PI3 kinase (PI3K), also known as VPS34, and class I PI3K regulatory subunit p85, in mice heart (Figure 6A and 6B, Figure S1G and S1H, and Figure S3). Consistent with normalized PIP levels, ALCAT1 deficiency or inhibition by Juve downregulated the expression of the two PI3Ks in the hearts of TAZ_KD_ and TAZ^cKO^ mice (Figure 6A and 6B, Figure S1G and S1H, and Figure S3). Moreover, the glycosylated form of lysosomal-associated membrane protein 1 (LAMP1), a regulator of lysosomal integrity, pH, and catabolic capacity^36^, was more abundant and likely post-translationally modified in TAZ-deficient hearts, as evidenced by the upward band shift on Western blot (Figure 6A and 6B, Figure S1G and S1H, and Figure S3). Ablation or inhibition of ALCAT1 suppressed the hyper-glycosylation of LAMP1 (Figure 6A and 6B, Figure S1G and S1H, and Figure S3), hinting at a restoration of lysosomal integrity and function in TAZ-deficient cells. In support of this notion, TAZ knockdown led to enlargement of lysosomes in mouse embryonic fibroblasts (MEFs) (Figure 6C and 6D), accompanied by an increase in lysosomal pH (Figure 6E). ALCAT1 deficiency or inhibition by Juve not only restored lysosome morphology to near-normal size but also normalized lysosomal pH (Figure 6C to 6E), suggesting a functional rescue of lysosomal activity.

**Figure 6.**
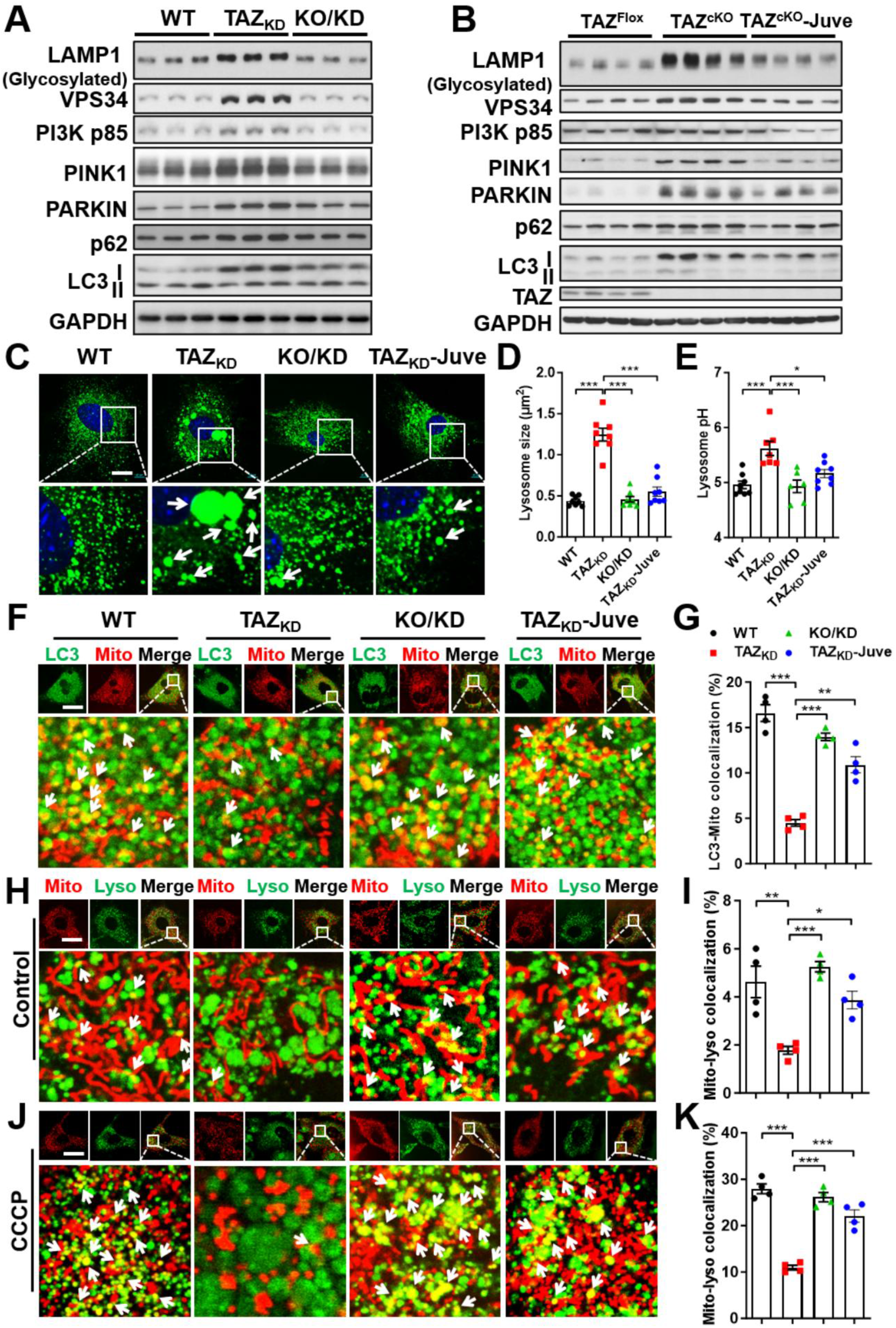
ALCAT1 deficiency or inhibition restores lysosome function and mitophagic flux in TAZ-deficient cells. **A-B**, Western blot analysis of the expression levels of proteins associated with autophagy, mitophagy, and lysosomal function in the hearts of WT, TAZ_KD_, and KO/KD mice (**A**, n=3), and the TAZ floxed and TAZ^cKO^ mice treated with vehicle or Juve (**B**, n = 4). **C**-**D**, Confocal imaging analysis (**C**) and quantitative analysis of lysosome size (**D**) in WT, TAZ_KD_ and KO/KD MEFs or TAZ_KD_ MEFs treated with Juve. Lysosomes and nuclei were stained by LysoTracker Green and Hoechst 33342, respectively. Arrows highlight enlarged lysosomes. n = 8. Scale bar, 20 μm. **E**, Lysosomal pH measurement in WT, TAZ_KD_, and KO/KD MEFs or TAZ_KD_ MEFs treated with Juve. n = 6-8. **F**-**G**, Confocal imaging analysis (**F**) and quantitative analysis (**G**) of the colocalization of mitochondria with autophagosomes in WT, TAZ_KD_, and KO/KD MEFs or TAZ_KD_ MEFs treated with Juve. MEFs were infected with GFP-LC3 adenovirus to label autophagosomes. Mitochondria were stained by MitoTracker Red. Cells were serum-starved in the presence of bafilomycin A1 to prevent autophagosome-lysosome fusion. Arrows highlight mitochondria colocalized with autophagosomes. n = 4. Scale bar, 40 μm.**H**-**I**, Confocal imaging analysis (**H**) and quantitative analysis (**I**) of the colocalization of mitochondria with lysosomes in WT, TAZ_KD_, and KO/KD MEFs or TAZ_KD_ MEFs treated with Juve under basal conditions. Mitochondria and lysosomes were stained by MitoTracker Red and LysoTracker Green, respectively. Arrows highlight mitochondria colocalized with lysosomes. n = 4. Scale bar, 40 μm. **J**-**K**, Confocal imaging analysis (**J**) and quantitative analysis (**K**) of the colocalization of mitochondria with lysosomes in WT, TAZ_KD_, and KO/KD MEFs or TAZ_KD_ MEFs treated with Juve in response to CCCP treatment. Mitochondria and lysosomes were stained by MitoTracker Red and LysoTracker Green, respectively, followed by treatment with CCCP (10 μM) for 1 hour to induce mitophagy. Arrows highlight mitochondria colocalized with lysosomes. n = 4. Scale bar, 40 μm. Data are presented as mean ± SEM. Statistical analysis was performed using one-way ANOVA with Tukey’s post hoc test. *p<0.01, **p<0.01, ***p<0.001.

To assess how ALCAT1 regulates defective mitophagy in BTHS, we monitored the distinct steps of mitophagic flux in MEFs. First, MEFs were infected with GFP-LC3 adenovirus to label autophagosomes and stained with MitoTracker red to label mitochondria. In the presence of bafilomycin A1 which prevents autophagosome-lysosome fusion, TAZ deficiency reduced the co-localization of mitochondria with autophagosomes in response to serum starvation (Figure 6F and 6G). In contrast, ALCAT1 deficiency or inhibition by Juve increased mitochondrial recruitment into autophagosomes (Figure 6F and 6G). Next, we determined the effect of ALCAT1 on autophagic consumption, a process that deliver damaged mitochondria to lysosomes. Under basal conditions, TAZ_KD_ MEFs exhibited less colocalization of mitochondria with lysosomes relative to WT MEFs, suggesting a defect in the lysosomal degradation of damaged mitochondria (Figure 6H and 6I). ALCAT1 deficiency or Juve treatment increased mitochondrial localization with lysosomes (Figure 6H and 6I). These findings were confirmed in cells treated with carbonyl cyanide m-chlorophenyl hydrazone (CCCP), a mitochondrial uncoupler that triggers mitophagy. CCCP treatment robustly induced mitophagy in WT MEFs (Figure 6J and 6K). However, TAZ_KD_ MEFs exhibited less mitochondria-lysosome colocalizations despite mitochondrial depolarization by CCCP (Figure 6J and 6K). Again, ALCAT1 deficiency or Juve treatment restored CCCP-induced mitophagy in TAZ-deficient MEFs versus WT controls (Figure 6J and 6K).

## Discussion

BTHS is a devastating mitochondrial disorder for which no effective disease-modifying therapy is available. Although defective CL remodeling caused by *Taz* mutations is the defining biochemical feature of BTHS, the mechanisms linking CL deficiency to mitochondrial dysfunction and cardiomyopathy remain incompletely understood. In this study, we identified ALCAT1 as an unappreciated mediator of mitochondria homeostasis in BTHS. ALCAT1 expression was markedly induced in TAZ-deficient hearts and its genetic ablation or pharmacological inhibition by Juve improved cardiac function, reduced myocardial fibrosis, and restored mitochondrial function in both whole-body knockdown and cardiac-specific TAZ knockout mice. Mechanistically, our findings revealed an ALCAT1-dependent PI-PIP-lysosome-mitophagy signaling axis that links TAZ deficiency to defective mitochondrial quality control.

A particularly surprising finding of this study was that the therapeutic effects of ALCAT1 inhibition occurred without correction of the canonical CL abnormalities. The prevailing view of BTHS pathogenesis emphasizes defective CL remodeling, depletion of mature TLCL, and accumulation of MLCL, acting to impair respiratory chain organization and mitochondrial energetics^8,37^. However, despite robust improvements in mitochondrial and cardiac function, neither *Alcat1* deletion nor Juve treatment restored the CL defects caused by TAZ deficiency, positing that recovery in clinical BTHS may not require restoration of CL. Although both TAZ and ALCAT1 participate in CL remodeling, they have distinct biochemical properties, subcellular localizations, and physiological effects. TAZ is primarily localized to the inner mitochondrial membrane, where it remodels CL through transacylation reactions using other phospholipids as acyl donors^7^. In contrast, ALCAT1 is enriched in mitochondria-associated membranes and remodels CL by reacylating lysocardiolipin^17^. We previously showed that ALCAT1 deficiency significantly increases TLCL levels by preventing CL peroxidation^18,24^. This effect may depend, at least in part, on increased TAZ expression, which is supported by our new finding that ALCAT1 ablation increased cardiac TAZ protein levels. This could explain why ALCAT1 deficiency or inhibition failed to restore TLCL level in TAZ deficient mice. These findings indicate that the benefits of ALCAT1 inhibition are mostly through a CL-independent mechanism and downstream of TAZ loss.

Another finding of this study is the pivotal role of dysregulated PI remodeling in BTHS pathogenesis. In addition to abnormalities in CL, TAZ deficiency caused marked accumulation of PI and multiple PIPs in the heart, along with increased VPS34 and the PI3K regulatory subunit p85. PIPs coordinate autophagosome biogenesis, lysosomal trafficking, membrane fusion, and cargo degradation^33–35^. Dysregulation of PIPs are implicated in lysosomal dysfunction and impaired autophagic clearance in aging and degenerative diseases^29,38^. TAZ deficiency resulted in the accumulation of autophagic vacuoles containing damaged mitochondria, together with lysosomal enlargement, increased lysosomal pH, and abnormal LAMP1 glycosylation, all indicative of lysosomal dysfunction^38,39^. These data suggest that excessive or dysregulated PI/PIP remodeling disrupts lysosomal function and mitophagic flux rather than promoting productive mitochondrial quality control in BTHS. In addition to remodeling CL, ALCAT1 catalyzes the pathological remodeling of phosphatidylglycerol and PI ^17,28^. As well, genetic ablation or pharmacological inhibition of ALCAT1 by Juve effectively normalized PI and PIP levels, suppressed VPS34 and PI3K activation, restored lysosomal morphology and acidification, and rescued mitophagic flux. These effects were accompanied by improvements in mitochondrial ultrastructure and respiratory function. These findings indicate that ALCAT1 is upstream of PIP dysregulation and define an ALCAT1-PI/PIP-lysosome-mitophagy axis that ties TAZ deficiency to defective mitochondrial quality control in BTHS.

Despite intensive efforts, no disease-modifying therapy has been approved for BTHS, and clinical management is limited to supportive treatment for heart failure, arrhythmias, and neutropenia^40^. For example, dietary supplementation with safflower oil-derived linoleic acid delayed the onset of cardiomyopathy in mice that mimicked BTHS, although its efficacy declined with disease progression^41^. Adeno-associated virus (AAV)-mediated TAZ replacement also showed efficacy in mouse models of BTHS^32,42^. However, moving this to the clinic will likely be challenging given the need to target end-organ selectively, potential immune responses to AAV capsid proteins and uncertainty regarding long-term transgene expression. Elamipretide (SS-31), a mitochondria-targeting CL-binding tetrapeptide, improved skeletal muscle strength in individuals with BTHS in the TAZPOWER phase 2/3 clinical trial^43,44^. However, improvements in cardiac function during the randomized crossover phase were modest^43–46^. Current therapeutic approaches to BTHS have largely focused on restoring TAZ function or CL homeostasis. Our findings provide an alternative strategy that targets downstream mitochondrial quality control, despite persistent CL abnormalities. Juve is a potent, first-in-class ALCAT1 inhibitor, with an IC_50_ of 3 nM, and a favorable safety and tolerability profile^24^. Juve has also shown therapeutic efficacy in preclinical models of cardiovascular and metabolic disease characterized by oxidative stress and mitochondrial dysfunction^24,25,47^. These findings support further development of Juve as a disease-modifying therapy for BTHS.

In summary, TAZ deficiency induces ALCAT1 expression, resulting in maladaptive PI remodeling and PIP dysregulation, lysosomal dysfunction and impaired mitophagy. This pathway amplifies the mitochondrial damage initiated by CL deficiency and contributes to progressive cardiac dysfunction. Genetic deletion of *ALCAT1* or pharmacological inhibition of ALCAT1 restored lysosomal function, mitophagic flux, mitochondrial homeostasis and cardiac performance in two mouse models of BTHS, despite persistent CL abnormalities. These findings identify a CL-independent pathogenic mechanism downstream of TAZ loss and provide preclinical proof of concept for ALCAT1 inhibition as a first-in-class disease-modifying strategy for BTHS and, potentially, other mitochondrial disorders characterized by defective mitophagy.

## Materials and Methods

### Reagents

Anti-PIK3C3/VPS34 (4263), PI3K p85 (4292), and LC3 (2775) antibodies were purchased from Cell Signaling Technology. Anti-LAMP1 (ab24170) and PARKIN (ab15954) antibodies were from Abcam. Anti-PINK1 (23274-1-AP) antibody was from Proteintech. Anti-SQSTM1/p62 (P0067) antibody was from MilliporeSigma. Anti-GAPDH (sc-32233) antibody was from Santa Cruz Biotechnology. Anti-TAZ antibody was a gift from Dr. Steven M. Claypool (Johns Hopkins University School of Medicine)^3^. ALCAT1 antibody was kindly provided by Dr. Hiroyuki Arai (University of Tokyo). HRP conjugated goat anti-rabbit IgG Fc secondary antibody (31463) and HRP conjugated goat anti-mouse IgG (H+L) secondary antibody (31430) were from Thermo Fisher Scientific. Carbonyl cyanide 3-chlorophenylhydrazone (CCCP; C2759), carbonyl cyanide 4-(trifluoromethoxy)phenylhydrazone (FCCP; C2920), oligomycin A (75351), rotenone (R8875), antimycin A (A8674), N,N,N’,N’-tetramethyl-p-phenylenediamine (TMPD; T7394), ascorbic acid (1043003), sodium azide (S8032), succinate (S9512), phosphatase inhibitor cocktail 2 (P5726), and phosphatase inhibitor cocktail 3 (P0044) were purchased from MilliporeSigma. EDTA-free protease inhibitor cocktail (11873580001) was from Roche. MitoTracker™ Red CMXRos (M7512), LysoTracker™ Green DND-26 (L7526), Lysosensor Yellow/Blue DND-160 (L7545) Hoechst 33342 solution (62249), TRIzol (15596018), SuperScript™ IV first-strand synthesis system (18091200), and SYBR™ green PCR master mix (4309155) were from Thermo Fisher Scientific. Doxycycline hyclate (446061000) was from Acros Organics.

### Animal care

C57BL/6 mice with targeted deletion of the ALCAT1 gene were generated as previously described^18^. B6.Cg-*Gt(ROSA)26Sor^tm^*^37*(H1/tetO-RNAi:Tafazzin)Arte*^/ZkhuJ mouse, a tetracycline inducible shRNA-mediated *Tafazzin* knockdown (TAZ_KD_) mouse model of BTHS^30^, was purchased from The Jackson Laboratory (Stock No: 014648). TAZ floxed mice were kindly provided by Dr. Xi Fang (University of California, San Diego)^12^. *Myh6-cre* mice were purchased from The Jackson Laboratory (Stock No: 011038). ALCAT1 knockout mice were cross-bred with TAZ_KD_ mice to generate TAZ_KD_ mice with target deletion of ALCAT1 (KO/KD). TAZ floxed mice were cross-bred with *Myh6-cre* mice to generate cardiac-specific TAZ knockout (TAZ^cKO^). All animals were maintained in environmentally controlled conditions with a 12 h light/12 h dark cycle and free access to food and water. WT, TAZ_KD_, and KO/KD mice were fed a diet containing doxycycline (625 mg/Kg) or had access to drinking water containing doxycycline (2 mg/mL) starting at 1-month of age and continued to the end of the experiments. Only male mice were used in experiments because BTHS is an X-linked disorder that predominantly affects males in human. Animals were randomly assigned to experimental groups when applicable. No animal or experimental data were excluded from the analyses. Experiments involving animals were approved by the Institutional Animal Care and Use Committee, and used a protocol in accordance with NIH guidelines (NIH publication no. 86-23 [1985]).

### Echocardiography

Echocardiographic analysis of mice hearts was performed by using a VisualSonics Vevo 2100 Imaging System and an MS550D transducer (VisualSonics, Toronto, ON, Canada). M-mode short axis images of the left ventricular (LV) were analyzed to measure the following parameters: diastolic and systolic interventricular septal wall thickness (IVSd and IVSs), diastolic and systolic LV internal dimension (LVIDd and LVIDs), diastolic and systolic LV posterior wall thickness (LVPWd and LVPWs), LV ejection fraction (EF), and LV fractional shortening (FS).

### Grip strength test

Forelimb grip strength was measured with a grip strength meter. Mice were allowed to grasp a horizontal metal grid and were gently pulled backward by the tail until they released the grid. The peak force generated immediately before release was recorded by the instrument. Each mouse was tested five consecutive times with at least 1 minute of rest between measurements. The grip strength test was conducted at the San Antonio Nathan Shock Center Aging Animal and Functional Assessment core.

### Treadmill exercise tolerance test

Exercise capacity was assessed using a motorized treadmill system. Mice were acclimated to the treadmill for 3 consecutive days before testing. On the day of the experiment, mice were subjected to a graded exercise protocol beginning at 5 m/min, with the running speed increased by 2 m/min every 2 min until exhaustion. Exhaustion was defined as the inability of the mouse to resume running after remaining on the shock grid for more than 10 s despite gentle encouragement. Total running distance and work output were recorded and analyzed as indicators of exercise tolerance. Work output (J) was calculated using the following equation: body weight (kg) × gravitational acceleration (9.8 m/s²) × vertical distance traveled (m), where vertical distance was determined from the treadmill running distance and incline angle. Treadmill exercise tolerance tests were conducted at the San Antonio Nathan Shock Center Aging Animal and Functional Assessment core.

### Histological analysis

Heart samples were isolated and fixed with 4% paraformaldehyde for 48 h. Fixed hearts were dehydrated and embedded in paraffin, and 5 μm sections were cut with a Leica RM-2162 (Leica, Bensheim, Germany). Hematoxylin and Eosin (H&E) and Masson’s trichrome staining were performed as previously described^27^. The size of cardiomyocytes and collagen volume fraction were quantified using ImageJ software (NIH).

### Transmission electron microscopy

The mitochondrial ultrastructure in mouse LV was evaluated by using transmission electron microscopy. Heart samples from the same site of LV were fixed in 5% glutaraldehyde and 4% paraformaldehyde in 0.1 M sodium cacodylate buffer (pH 7.4) with 0.05% CaCl_2_ for 24 h. After washing in 0.1 M sodium cacodylate buffer, tissues were fixed overnight in 0.1 M cacodylate buffer containing 1% OsO_4_, dehydrated, and embedded in EMbed-812 resin (Electron Microscopy Sciences, 14121). The sections were stained with 2% uranyl acetate, followed by 0.4% lead citrate, and viewed with a JEOL JEM-2200FS 200kV electron microscope (Electron Microscopy Sciences Core at the UT Health San Antonio).

### Isolation and culture of adult mouse cardiomyocytes

Mice were injected with 8,000 IU/kg heparin sulfate and euthanized by isoflurane overdose. The aorta was exposed surgically and clamped, and the heart perfused with 3% collagenase type II in Tyrode’s solution (138 mM NaCl, 5.33 mM KCl, 0.33 mM NaH_2_PO_4_, 1.18 mM MgCl_2_, 10 mM HEPES, 10 mM taurine, and 10 mM glucose, pH 7.4) at 37 °C for 10 min. The ventricular apex was cut and transferred to Tyrode’s solution supplemented with 0.2% BSA. The tissue samples were cut into small pieces, dispersed by repeated pipetting, and filtered with a 220 μm mesh strainer. The myocytes were centrifuged at 500 g for 5 min and transferred to an uncoated 35 mm dish followed by calcium reintroduction. The myocytes were centrifuged and then cultured in minimum essential medium.

### Isolation and culture of mouse embryonic fibroblasts (MEFs)

MEFs were isolated at E12.5-13.5 from the time of copulation of female transgenic B6.Cg-*Gt(ROSA)26Sor^tm37(H1/tetO-RNAi:Tafazzin)Arte^*/ZkhuJ mice bred with male C57BL/6J mice. Each embryo was separated onto a dish with Hank’s balanced salt solution (Gibco, 14175). Head, limbs, and internal organs were removed. The head was collected and genomic DNA was isolated for genotyping using primers and PCR conditions previously described^30^. The remaining embryo was minced with a scalpel or razor blade and transferred to a 15 mL tube with 4 mL of collagenase solution (2 mg/mL collagenase IV [MilliporeSigma, C5138], 0.7 mg/mL DNase I [MilliporeSigma, D5025] and 10 mg/mL hyaluronidase [MilliporeSigma, H6254]), followed by incubation for 30–60 min in a 37°C until tissue chunks were absent. The solution was filtered through 100 μm mesh and washed with culture medium before allocating into culture dishes. MEFs were cultured in Dulbecco’s Modified Eagle’s Medium (MilliporeSigma, D5796) supplemented with 10% fetal bovine serum (Atlanta Biologicals, S11550H), 1x nonessential amino acids (Gibco, 11140-050), 1x essential amino acids (Gibco, 11130-051), 100 U/mL penicillin-streptomycin (Corning, 30-002-Cl), and 1.25 mg/mL amphotericin B (Corning, 30-003-CF). MEFs were cultured in completed medium containing doxycycline (1 μg/mL) for 3 days to induce *Taz* knockdown.

### Mitochondrial Oxygen consumption rate (OCR) and ROS production measurements

In cultured cardiomyocytes, mitochondrial OCR was characterized using a Seahorse XF24 analyzer (Seahorse Bioscience, North Billerica, MA, USA) at baseline, followed by exposure to 1 μM oligomycin (ATP synthase inhibitor), 1.5 μM FCCP (oxidative phosphorylation uncoupler), and a mixture of 1 μM rotenone (respiratory complex I inhibitor) with 1 μM Antimycin A (respiratory complex III inhibitor). After the measurements, the cells were lysed and protein concentrations were measured by using a BCA protein assay kit (Thermo Fisher Scientific, 23225). The OCR was normalized with the protein in each well. OCR in isolated mitochondria from fresh or frozen heart samples was analyzed as previously described^48^. Mitochondria (4 μg protein/well) isolated from fresh or frozen heart samples was loaded onto XF24 microplate. OCR in mitochondria isolated from fresh hearts was measured with succinate (10mM) as initial substrate, followed by ADP (4 mM), oligomycin (2.5ug/mL), FCCP (4uM), and Antimycin A (4uM). OCR in mitochondria isolated from frozen hearts was measured in response to a mixture of succinate (5 mM) and rotenone (2 μM), antimycin A (4 μM), a mixture of TMPD (0.5 mM) and ascorbate (1 mM), and sodium azide (50 mM). Real-time OCR was recorded two times during each conditional cycle. The ROS generation of isolated mitochondria from fresh heart tissues was detected indirectly by quantitatively measuring H_2_O_2_ production, as previously described^18^.

### Quantitative real time-PCR (qRT-PCR) analysis

Total RNA was extracted from hearts using TRIzol reagent, and RNA quantity was determined at 260 nm (NanoDrop 1000 Spectrometer, Thermal Scientific). Single-stranded cDNA was synthesized using the SuperScript IV reverse transcriptase system, and gene expression was determined by qRT-PCR performed using SYBR green PCR master mix and the 7300 Real-Time PCR System (Applied Biosystems, Waltham, MA, USA). The mRNA levels were determined using a standard curve method and normalized to the level of *Gapdh*. The primers used in PCR were:

*Gapdh*: AATGGTGAAGGTCGGTGTG, GTGGAGTCATACTGGAACATGTAG

*Anf*: GTGTACAGTGCGGTGTCCAA, ACCTCATCTTCTACCGGATC

*Bnp*: GCTGCTTTGGGCACAAGATAG, GGAGCTCTTCCTACAACAACTT

*β-Mhc*: AGGGCGACCTCAACGAGAT, CAGCAGACTCTGGAGGCTCTT

*Acta1*: GTTCGCGCTCTCTCTCCTCA, GCAACCACAGCATTGTC

*Collagen I*: GAGCGGAGAGTACTGGATCG, GTTCGGGCTGATGTACCAGT

*Collagen III*: ACCAAAAGGTGATGCTGGAC, GACCTCGTGCTCCAGTTAGC

### mtDNA copy number assays

Genomic DNA was extracted using the Wizard® Genomic DNA Purification Kit (A1125, Promega). mtDNA copy number was determined by PCR using mitochondrion-encoded NADH dehydrogenase 1 (Nd1) as the mtDNA markers and cyclophilin A as the nuclear DNA marker. The primers used were:

*Cyclophilin A*: ACACGCCATAATGGCACTCC, CAGTCTTGGCAGTGCAGAT

*Nd1*: GTGTACAGTGCGGTGTCCAA, ACCTCATCTTCTACCGGATC

### Lysosomal pH measurement

Lysosomal pH was determined using dextran conjugated Lysosensor Yellow/Blue DND-160. MEFs were seeded on a clear bottom 96-well-plate with black walls and incubated with 10 μM lysosensor for 10 min at 37℃ with 5% CO_2_. Cells were washed 3X in HBSS and the fluorescence was detected using a microplate reader with an excitation of 340 nm and an emission wavelength of 430 and 535 nm. To generate a pH calibration curve, cells were pre-incubated with 10 μM lysosensor for 10 min and then treated for 30 min with lysosomal equilibration buffers (10 μM monensin, 10 μM nigericin, 5 mM NaCl, 115 mM KCl, 1.3 mM MgSO_4_, and 25 mM MES, pH from 3.5 to 7.0). The ratio of emission 430/535 nm was then calculated for each sample. The pH values were determined from the linear standard curve generated via the pH calibration samples.

### Western blot analysis

Protein was separated by SDS-PAGE, and then transferred to PVDF membranes. After blocking in 5% non-fat milk, membranes were incubated with primary antibodies. The membranes were washed in TBST buffer and then incubated with horseradish peroxidase-labeled secondary antibodies and chemiluminescent substrate. Proteins in the immunoblot were visualized by exposing membranes to x-ray film.

### Lipidomics

The lipidomic analysis was carried out as previously described^18,49^. Total lipids from murine heart samples were analyzed with a triple-quadruple mass spectrometer (Thermo Electron TSQ Quantum Ultra, Trzin, Slovenia) controlled by Xcalibur (Thermo Fisher Scientific) system software. All the mass spectrometer spectra and tandem mass spectrometer spectra were acquired automatically by a customized sequence subroutine operated with Xcalibur software.

### Confocal imaging analysis

MEFs were stained with MitoTracker Red CMXRos (50 nM), LysoTracker Green DND-26 (100 nM), and Hoechst 33342 (1 μM) to visualize mitochondria, lysosomes, and nucleus, respectively. MEFs were infected with GFP-LP3 adenovirus to label autophagosomes. Cells were then serum-starved in the presence of bafilomycin A1 (1 nM) to prevent autophagosome-lysosome fusion. To induce mitophagy, MEFs were treated with CCCP (10 μM) for 1 hour in complete medium. Mitochondria were stained with MitoTracker Red CMXRos before CCCP treatment. In some studies, the cells were pre-treated with Juve (10 μ) for 24 hours before other treatments. Images were captured using a Leica Stellaris 5 confocal microscope (Leica Camera AG, Wetzlar, Germany) with a 63x/1.4 NA oil immersion objective. Multiple fields of view were selected at random and imaged for analysis with the Leica Application Suite X or ImageJ software (NIH).

### Statistical analysis

Data were represented as mean ± SEM. Statistical analysis was performed using one-way ANOVA with Tukey’s post hoc test or unpaired Student t-test using GraphPad Prism 10. Biological significance was considered statistically significant at p < 0.05. *p < 0.05; **p < 0.01; ***p < 0.001.

## Acknowledgements

We would like to thank Dr. Steven M. Claypool at the Johns Hopkins University School of Medicine for providing us with the anti-TAZ antibody, Dr. Xianlin Han at UT Health San Antonio for technical help with lipidomic analysis, and Dr. Jeffrey S. Isenberg at City of Hope for reviewing and critical comments on the manuscript.

## Author contributions

Y.S. conceived and supervised the study. Y.S. and J.Z. designed the experiments. J.Z. performed the experiments, collected and analyzed the data. Y.Q. assisted on animal care and experiments. X.F. provided the TAZ floxed mice. W.H. gave suggestions on the experiment design. J.Z. and Y.S. wrote the manuscript. All authors read, reviewed, and approved of the final manuscript.

## Novelty and Significance

### What Is Known?

- Barth syndrome (BTHS) is caused by loss-of-function mutations in *TAFAZZIN* (*TAZ*), resulting in defective cardiolipin (CL) remodeling, mitochondrial dysfunction, and cardiomyopathy.
- Defective mitochondrial quality control and impaired mitophagy contribute to BTHS cardiomyopathy, but the underlying mechanisms remain incompletely understood.
- There are currently no disease-modifying therapies for BTHS, and therapeutic approaches have largely focused on restoring TAZ function or CL homeostasis.

### What New Information Does This Article Contribute?

- Genetic deletion or pharmacological inhibition of ALCAT1 restores mitochondrial function and ameliorates cardiomyopathy in two mouse models of BTHS despite persistent CL abnormalities.
- Upregulated ALCAT1 by TAZ deficiency caused pathological phosphatidylinositol remodeling that disrupts lysosomal function, mitophagic flux and mitocondrial quanlity control.
- Pharmacological inhibition of ALCAT1 with Juvenatin improves mitochondrial and cardiac function, establishing ALCAT1 as a promising disease-modifying therapeutic target for BTHS.

## Novelty and Significance

BTHS is a mitochondrial disorder defined by defective CL remodeling, and therapeutic strategies have therefore largely focused on restoring TAZ function or CL homeostasis. This study identifies ALCAT1 as a previously unrecognized mediator of BTHS cardiomyopathy and reveals a CL-independent mechanism linking TAZ deficiency to defective mitochondrial quality control. TAZ deficiency induces ALCAT1 and promotes aberrant phosphatidylinositol remodeling and phosphoinositide accumulation, leading to lysosomal dysfunction and impaired mitophagic flux. Genetic deletion or pharmacological inhibition of ALCAT1 by Juvenatin, a highly selective small molecule ALCAT1 inhibitor, restores lysosomal function, mitophagy, mitochondrial respiration, and cardiac performance without correcting the underlying CL abnormalities. Importantly, treatment with Juvenatin improves mitochondrial and cardiac function in two complementary mouse models of BTHS, including mice with established cardiomyopathy. These findings expand the current understanding of BTHS pathogenesis beyond CL deficiency and provide preclinical evidence that targeting downstream mitochondrial quality control through ALCAT1 inhibition may represent a disease-modifying therapeutic strategy for BTHS.

---

**Figure S1.**
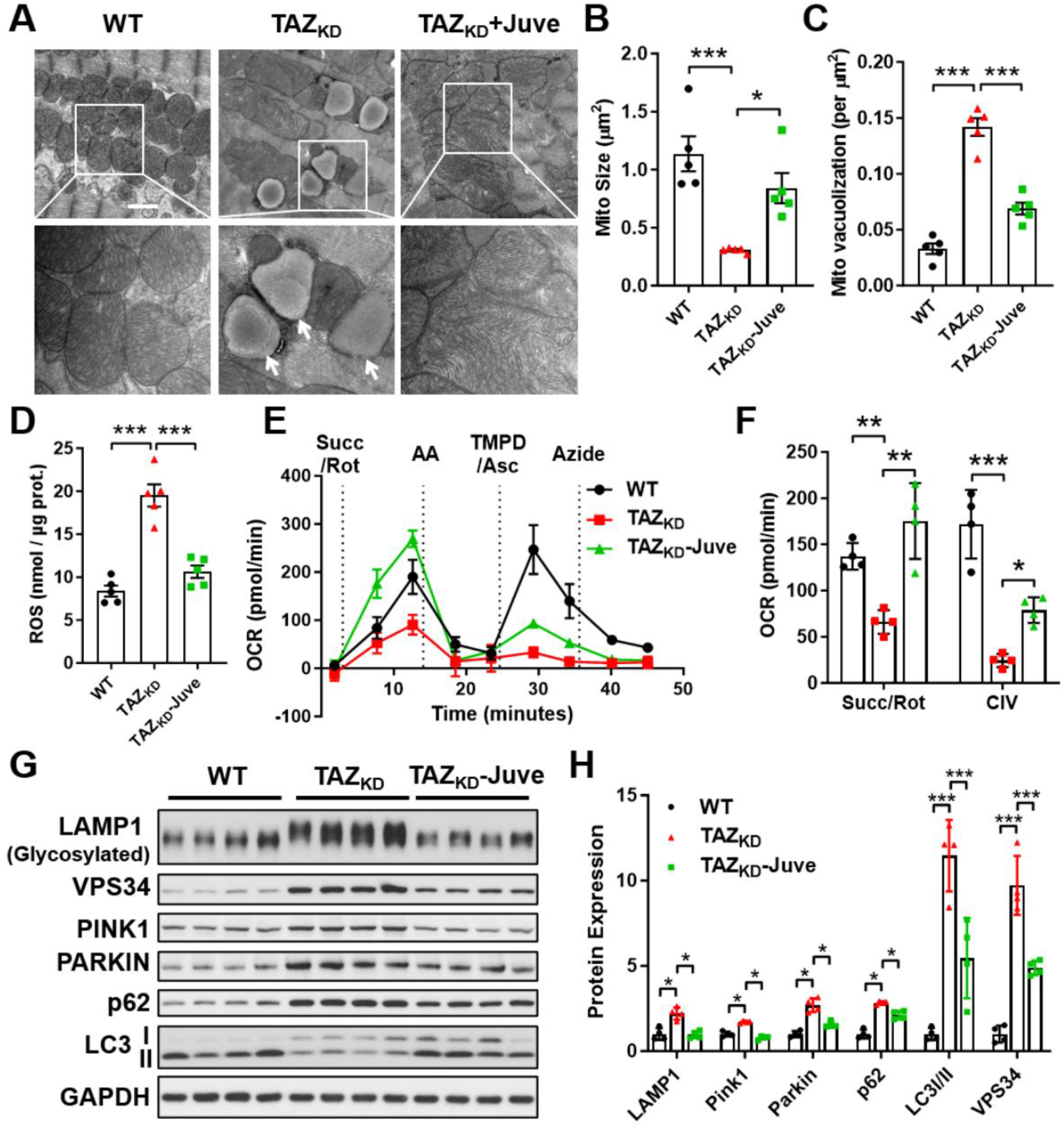
Inhibition of ALCAT1 by Juve restores mitochondrial function and mitophagy in TAZ_KD_ mice. **A-C**, TEM analysis of mitochondria morphology (**A**) and quantitative analysis of mitochondrial size (**B**) and vacuolization rate (**C**) in the hearts of WT and TAZ_KD_ mice treated with vehicle or Juve. n = 5. Scale bar, 2 μm. **D**, ROS production in isolated mitochondria from WT and TAZ_KD_ murine hearts treated with vehicle or Juve. n = 5. **E**-**F**, Seahorse analysis (**E**) and quantitative analysis (**F**) of mitochondrial OCR in mitochondria isolated from frozen hearts of WT and TAZ_KD_ mice treated with vehicle or Juve. n = 4. **G**-**H**, Western blot (**G**) and quantitative analysis (**H**) of the expression levels of proteins associated with autophagy, mitophagy, and lysosomal function in the hearts of WT and TAZ_KD_ mice treated with vehicle or Juve. n = 4. Data are presented as mean ± SEM. Statistical analysis was performed using one-way ANOVA with Tukey’s post hoc test. *p<0.01, **p<0.01, ***p<0.001.

**Figure S2.**
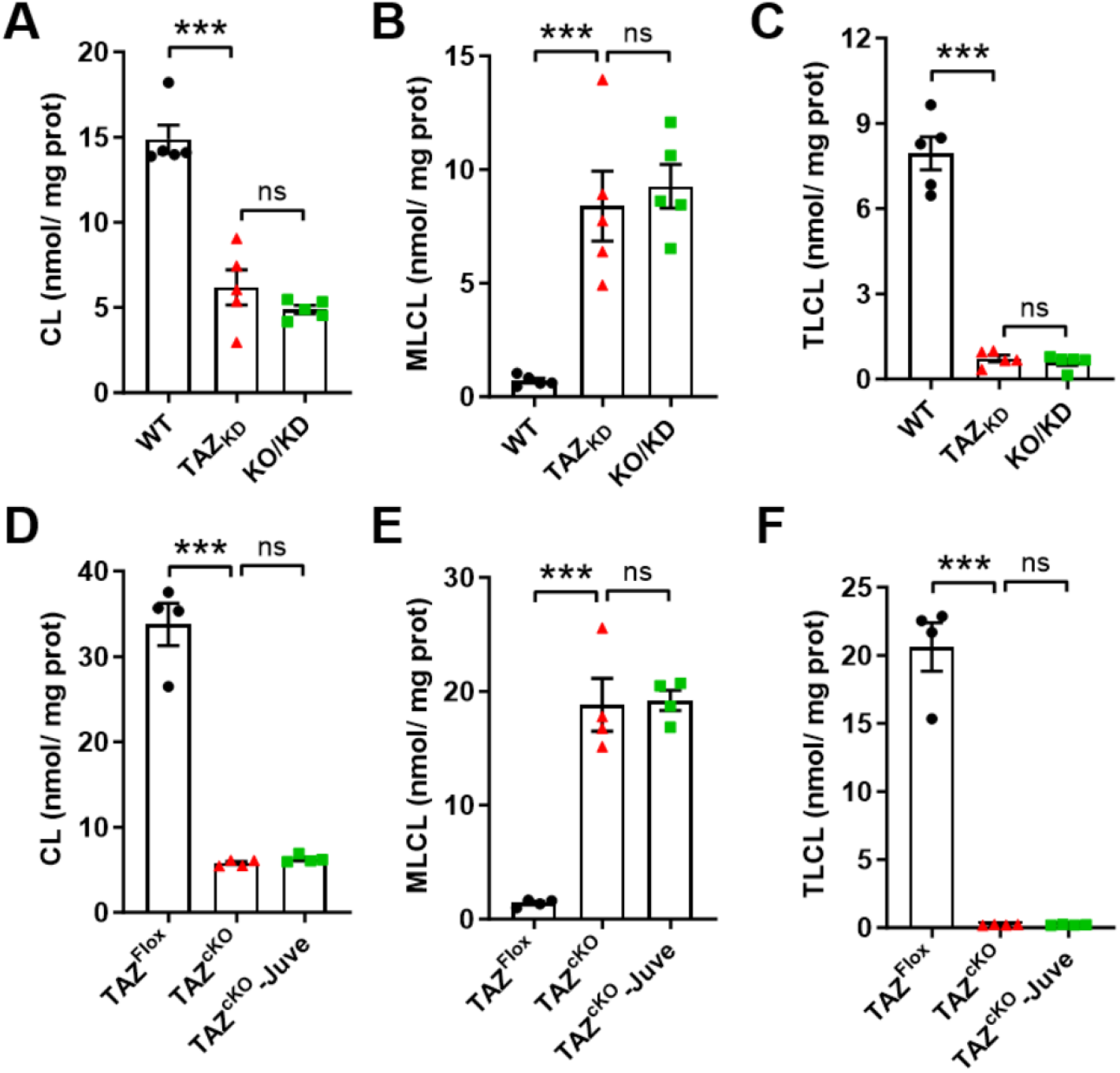
ALCAT1 deficiency or inhibition does not restore cardiolipin deficiency in TAZ-deficient hearts. **A-C**, Lipidomic analysis of CL (**A**), MLCL (**B**), and TLCL (**C**) levels in the hearts of WT, TAZ_KD_, and KO/KD mice. n = 5. **D**-**F**, Lipidomic analysis of CL (**D**), MLCL (**E**), and TLCL (**F**) levels in the heart of TAZ floxed and TAZ^cKO^ mice treated with vehicle or Juve. n = 4. Data are presented as mean ± SEM. Statistical analysis was performed using one-way ANOVA with Tukey’s post hoc test. ***p<0.001; ns, no significance.

**Figure S3.**
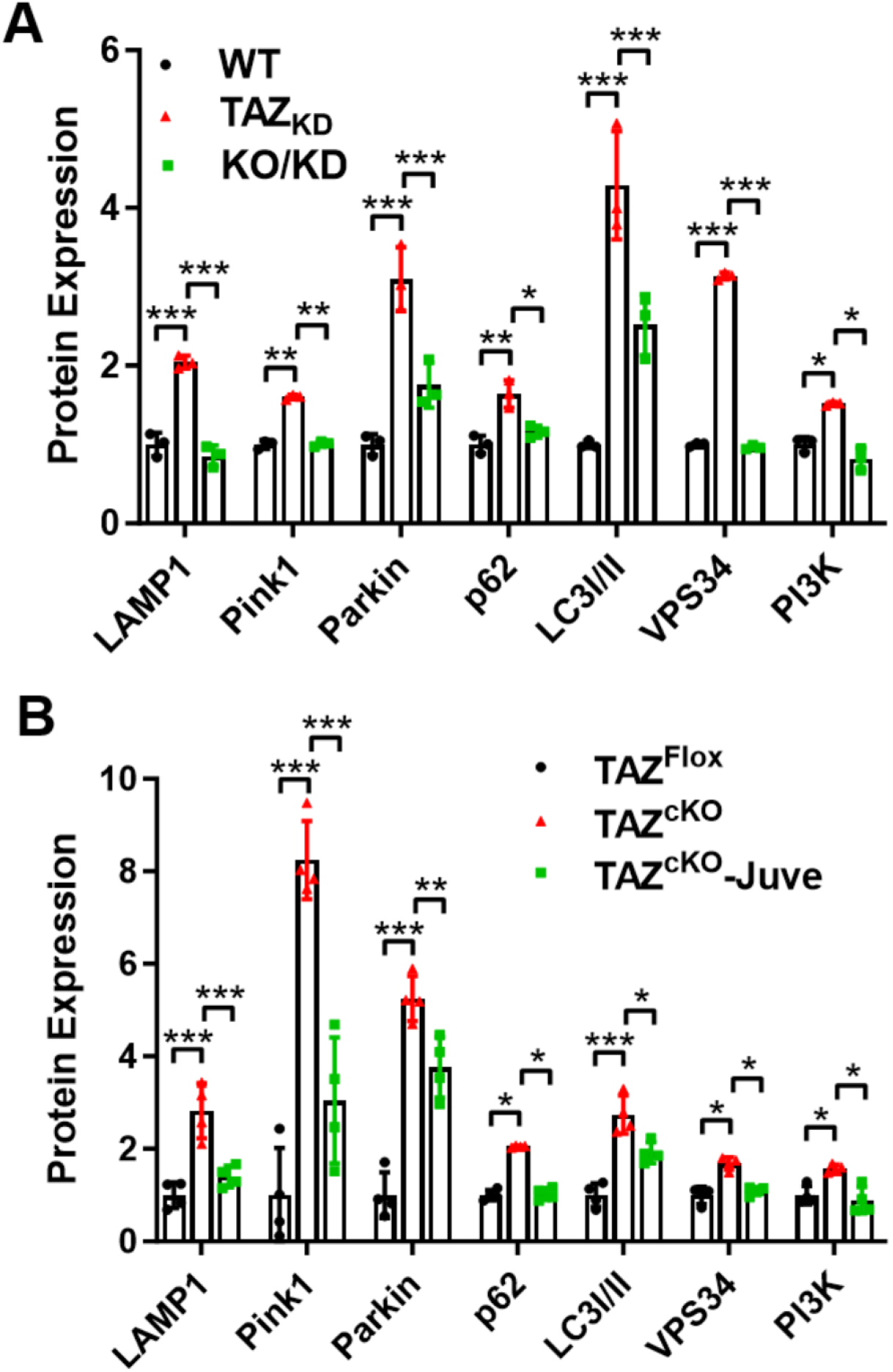
ALCAT1 deficiency or inhibition restore the expression of autophagy, mitophagy, and lysosomal function biomarkers in TAZ-deficient heart. **A**-**B**, Quantitative analysis of the expression levels of proteins associated with autophagy, mitophagy, and lysosomal function in the heart of WT, TAZ_KD_, and KO/KD mice (**A**, n=3) and the TAZ floxed and TAZ^cKO^ mice treated with vehicle or Juve (**B**, n = 4). Data are presented as mean ± SEM. Statistical analysis was performed using one-way ANOVA with Tukey’s post hoc test. *p<0.01, **p<0.01, ***p<0.001.

